# A microfluidic biodisplay

**DOI:** 10.1101/112110

**Authors:** Francesca Volpetti, Ekaterina Petrova, Sebastian J. Maerkl

## Abstract

Synthetically engineered cells are powerful and potentially useful biosensors, but it remains problematic to deploy such systems due to practical difficulties and biosafety concerns. To overcome these hurdles, we developed a microfluidic device that serves as an interface between an engineered cellular system, environment, and user. We created a biodisplay consisting of 768 individually programmable biopixels and demonstrated that it can perform multiplexed, continuous sampling. The biodisplay detected 10 µg/l sodium-arsenite in tap water using a research grade fluorescent microscope, and reported arsenic contamination down to 20 µg/l with an easy to interpret “skull and crossbones” symbol detectable with a low-cost USB microscope or by eye. The biodisplay was designed to prevent release of chemical or biological material to avoid environmental contamination. The microfluidic biodisplay thus provides a practical solution for the deployment and application of engineered cellular systems.

## GRAPHICAL ABSTRACT

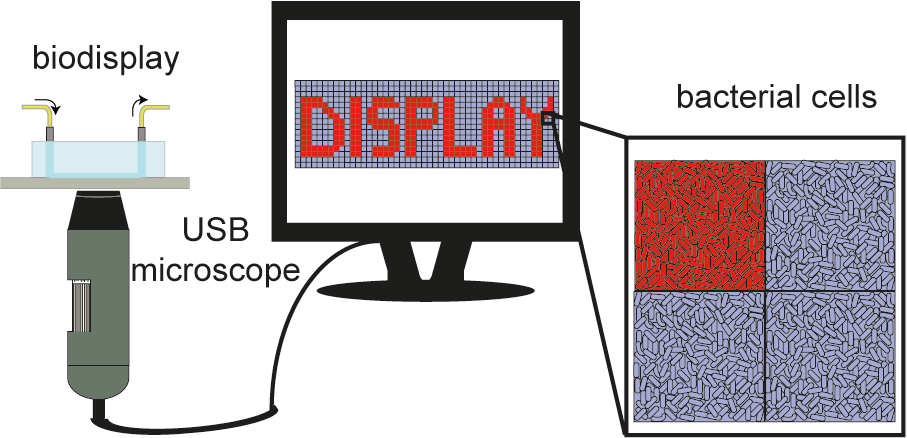

With the advent of synthetic biology, a number of biological sensors have been engineered capable of monitoring the environment^1,2^. Although such synthetic biological systems are in principle powerful sensors and information processing units, they often lack direct applicability because suitable interfaces between the environment, the user and the engineered biological system don’t exist^3,4^. The lack of such interfaces results in problems related to safely deploying genetically modified organisms (GMOs), while making it possible for them to interact with the environment and user^5^. Aside from safety concerns, interfaces can solve practical problems such as keeping the engineered biological system alive for extended periods of time, automatic and frequent sampling of the environment, and facilitating readout.

Microfluidics originated as a tool to enable analytical measurements in chemistry and biology^6^. In the last decades, microfluidic devices have found a plethora of applications in biology^6^ spanning high-throughput screening^7^, cell-based assays^8,9^, and molecular diagnostics^10^. The use of microfluidic devices increases throughput, reduces cost, and can enable novel measurements^11^. In most instances the purpose of microfluidics is to enable or conduct analytical measurements of molecules or cells. Few examples depart from this dogmatic application of microfluidics and instead employ microfluidic devices as soft robots capable of movement and camouflage^12^, or as microfluidic games^13^. One recent example demonstrated how engineered biological systems can be deployed by combining bacterial sensors with passive microfluidic channels in a wearable hydrogel-elastomer hybrid device capable of sensing and reporting the presence of diacetylphlorogluicinol (DAPG), isopropyl β-d-1thiogalactopyranoside (IPTG), acyl homoserine lactones (AHL) and rhamnose^14^.

We previously developed a method to culture and interrogate 1’152 *S. cerevisiae* strains on a microfluidic device. Each strain was cultured in a dedicated micro-chemostat and was interrogated optically to provide single-cell, phenotypic information on each strain. We applied this platform to the comprehensive, singlecell analysis of yeast proteome dynamics^15^ and a detailed characterization and modeling of transcriptional regulation in yeast^16^. Here we show that this approach can be used to generate a bacterial biodisplay. Unlike previous bacterial biodisplays^17^, each of the 768 pixels can be independently programmed with a specific bacterial strain or clone. We used the biodisplay to characterize the response of several engineered bacterial strains to small molecule analytes for extended periods of time. To demonstrate environmental monitoring, we programmed the biodisplay with a “skull and crossbones” symbol using a bacterial arsenic sensor strain. The symbol is displayed when arsenic contaminated water is sensed by the biodisplay. We also demonstrated a multiplexed biodisplay that reports on the presence or absence of two small molecules: arabinose and arsenic. The use of an easy-to-interpret symbolic display drastically simplifies analysis, and is readily understood by a layman without any special hardware or analysis requirements. To enable field-deployment of our biodisplay we showed that readout can be conducted using an off-the-shelf low-cost USB microscope, a cellphone, or direct visual inspection. Finally, to increase shelf-life and facilitate shipping / deployment of the biodisplay we programmed the device with *B. subtilis* spores, which allowed storage of the display at 80ºC for 1 month without any detectable decrease in viability or sensing functionality.

## RESULTS AND DISCUSSION

### Biodisplay programming, culturing, sampling and readout

The biodisplay has a resolution of 48 × 16 for a total of 768 programmable pixels with a density of 64 pixels per inch (Figure 1a). Each pixel consists of a 250 μm square cell chamber, two side channels for medium and sample introduction, sandwich valves to isolate adjacent pixels and a neck valve to isolate the pixel from the side channels. Each pixel of the biodisplay can be programmed with a different bacterial strain. Standard microarray spotting was used to array bacterial cells or spores on an epoxy-coated glass slide. *E. coli* was spotted in medium containing 10% glycerol to preserve cell viability, as we observed that dried *E. coli* spots failed to regrow after spotting. *B. subtilis* spores could be spotted in water without addition of glycerol. The bacterial array was then aligned to the microfluidic chip and bonded at 37°C for 1 hour.

**Figure 1.**
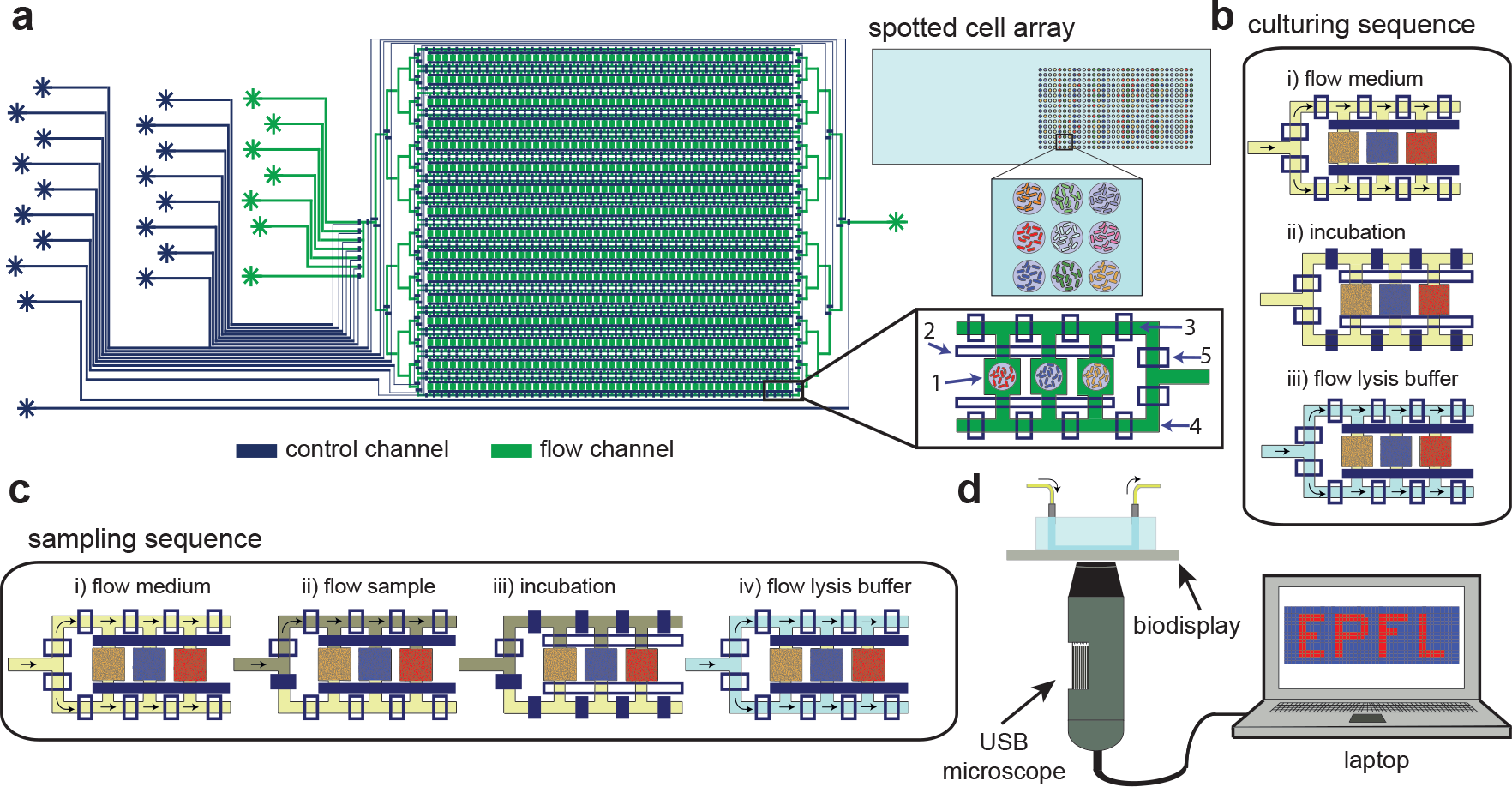
Biodisplay schematic and use. (**a**) Design of the two-layer microfluidic device (control channels in blue and flow channels in green). The device is composed of 16 rows and 48 columns for a total of 768 biopixels. Each pixel consists of a chamber (1), chamber valve (2), sandwich valves (3) and side channels (4). The side channels in turn can be specifically addressed using the side channel valves (5). A bacterial cell array is spotted on a glass slide and aligned with the PDMS chip. (**b**) The culturing sequence consists of 3 steps: i) flowing medium with the chambers closed (10 minutes), ii) incubation with the sandwich valves closed and chamber valves open (45 minutes), and iii) flowing lysis buffer with the chamber closed (10 minutes). (**c**) The sampling sequence consists of 4 steps: i) flowing the medium in both side channels (10 minutes), ii) flowing the sample in one side channel (10 minutes), iii) incubation with both medium and sample (45 minutes), and iv) flowing lysis buffer (10 minutes). The sequence is than repeated continuously for the extent of the experiment. (**d**) The biodisplay can be imaged using a standard research grade epifluorescent microscope, a USB fluorescent microscope, a cellphone camera, or by eye.

After assembling and bonding the biodisplay, cells were allowed to recover overnight using an automated culturing routine (Figure 1b). A medium reservoir was attached to the device and the waste outlet was connected to a receptacle containing 70% aq. EtOH to immediately sterilize and contain the device outflow. First, the entire device was loaded with medium by outgas priming^18^. Once the device was fully primed with medium, cells were cultured with the following three-step culturing routine: i) flowing medium through the side channels (10 minutes), ii) allowing medium to diffuse into the biopixels (45 minutes) and iii) flowing lysis buffer through the side channels (10 minutes). Whenever fluid was flowed through the side channels, the biopixel chambers were sealed off from the side channels using a set of valves. In step (ii) the biopixel valves were opened to allow diffusion of molecules in and out of the biopixel chamber. During this step, individual pixels were isolated from one another by closed sandwich valves. The lysis segment of the culturing routine prevented biofilm and microcolony formation outside of the biopixels^19^. By actively preventing accumulation and growth of cells in areas other than the biopixels we avoided device clogging and allowed experiments to proceed for several days; the longest experiment conducted here lasted one week. Culturing was performed at 37ºC using a thermo glass plate (Okolab). Culturing at 37ºC increased growth rate, leading to faster accumulation of biomass, but was otherwise not required and incubation at room temperature was also possible. After overnight culturing, each bacterial spot had outgrown, and populated its dedicated biopixel. We did not observe any contamination resulting from microarraying. Cells occasionally populated empty biopixels during culturing, which was avoided by spotting a non-fluorescent *E. coli* strain in otherwise empty biopixels.

Once the biodisplay was online, we switched from the culturing routine to a sampling routine that enabled frequent testing of a liquid sample (Figure 1c). The sampling routine was identical to the culturing routine, with an additional step added prior to diffusive mixing of the pixel and side channels. After flowing medium through both side channels, the sample was flowed through one of the side channels. During the diffusion step, both medium and sample were allowed to diffuse into the biopixel chambers. This process allowed us to aspirate a water sample directly, rather than having to premix it manually with the medium solution. By using a dedicated sampling port, it was possible to automatically sample a source for extended periods of time in 75 minute intervals without user interference.

We employed a standard research grade epi-fluorescent microscope (Nikon) and a handheld, low-cost USB fluorescent microscope (Dino-Lite) for device readout and quantitation (Figure 1d). The standard epifluorescent microscope equipped with a motorized stage was programmed to image each biopixel individually whereas the handheld USB microscope could image either the entire display at once or acquire several sub-sections. Both approaches were used to acquire time-course and endpoint measurements. We also demonstrated that the biodisplay could be observed directly by eye using a LED flashlight and an emission filter, and images could be acquired using a standard cellphone camera.

### Multiplexed characterization of bacterial strains

We tested the biodisplay device by culturing and characterizing nine *E. coli* strains engineered to induce GFP or RFP expression in response to arabinose (Table S1). These strains were obtained from the iGEM registry of standard biological parts and were generated by various student teams. The fluorescent protein genes were under the regulation of a pBAD promoter or derivatives thereof. We programmed the biodisplay with the nine strains, each strain filling 2 columns of the device for a total of 32 pixels per strain. Strains were induced with arabinose after overnight culturing at 37°C. Consequent culturing and induction were performed at room temperature (23-24ºC) (Figure 2a) or 37ºC (Figure S1). Room temperature induction was used in order to assess the performance of the biodisplay and cell-based sensors at ambient temperature in order to reduce platform complexity by eliminating the need for a temperature control element.

**Figure 2.**
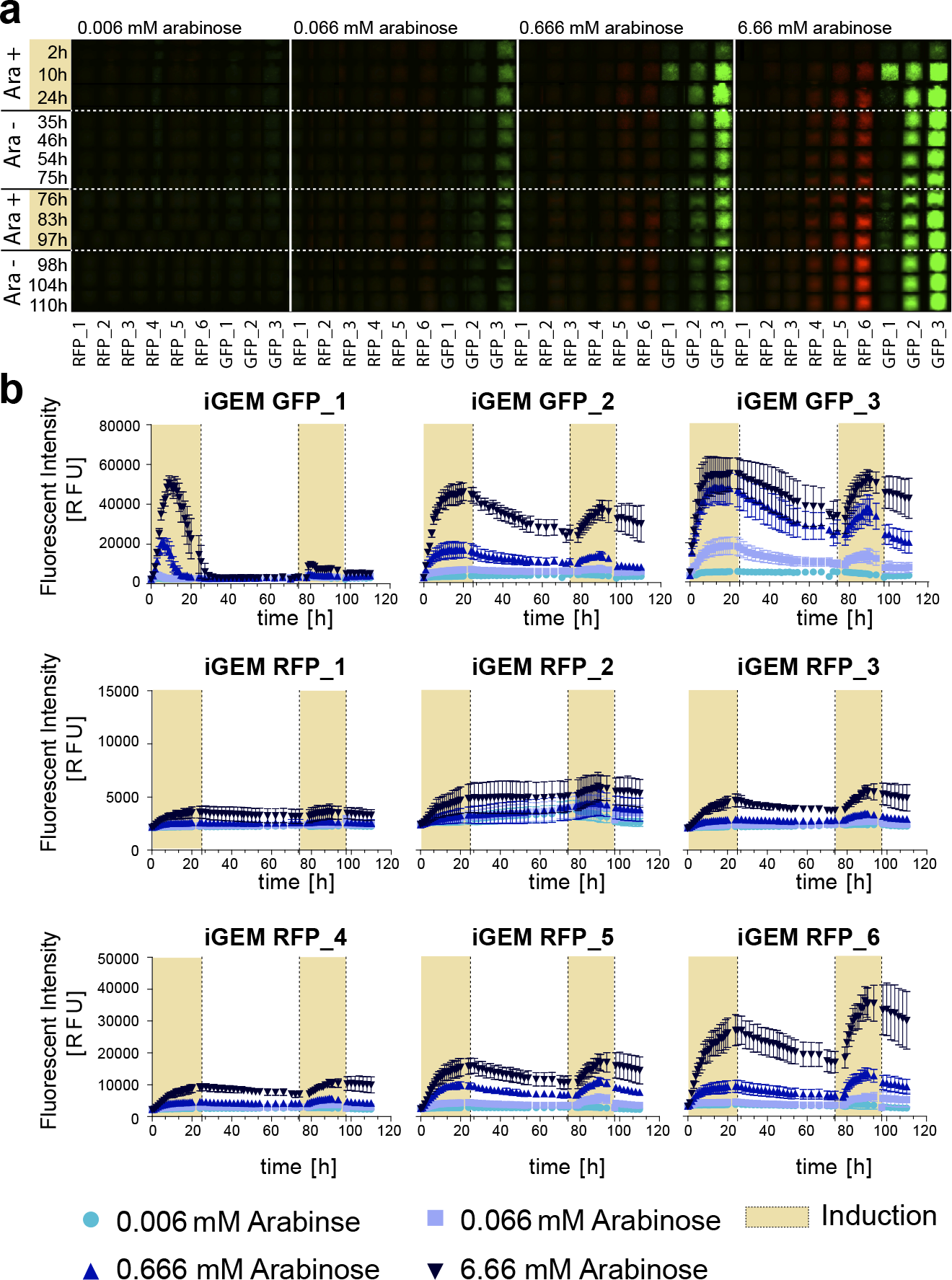
Using the biodisplay for strain characterization. (**a**) Fluorescence kymograms of nine arabinose sensing *E. coli* strains. Shown is a representative biopixel for each strain and condition tested. The strains were induced twice for 24 hours with 0.006 mM, 0.066 mM, 0.666 mM and 6.66 mM arabinose. (**b**) Quantitative analysis of fluorescence pixel intensities for each strain under four arabinose concentrations. The yellow background denotes the two induction periods. Each data point represents the average of eight pixels with error bars representing to the std. deviation of the means.

We designed the biodisplay device so that it was possible to specifically address four sets of four rows with a different solution. We used this modality to characterize each strain against four arabinose concentrations: 0.006 mM, 0.066 mM, 0.666 mM, 6.66 mM (0.0001%, 0.001%, 0.01% and 0.1% w/v) arabinose in lysogeny broth (LB) medium. Altogether, we tested 36 strain – inducer combinations by measuring nine strains in response to four arabinose concentrations with eight technical repeats for each combination. The biodisplay was placed on an automated research-grade fluorescent microscope and imaged over time.

We performed two consecutive 24 hour inductions, with the second induction starting 51 hours after the end of the first (Figure 2). Upon induction we observed a measurable signal in all strains. All three of the GFP strains showed a strong response to 6.66 mM arabinose while only strain #3 induced highly in response to 0.666 mM. Strains #1 and #2 induced to low levels in response to 0.666 mM and failed to induce to appreciable levels at lower arabinose concentrations, whereas strain #3 also induced in the presence of 0.066 mM arabinose. These results indicate that strains #2 and #3 could be used in combination to distinguish three different inducer concentrations ranging over 3 orders of magnitude. All RFP strains induced expression to differing degrees in response to 6.66 mM arabinose and in the cases of RFP strains #2-6 also to 0.666 mM arabinose. All strains, with the exception of strain GFP_1 slowly decreased in intensity during the 51 hour de-induction period. Strain GFP_1 exhibited a transient expression profile with signal beginning to decrease during the induction phase. GFP_1 is also the only strain that failed to induce during the second induction when cultured at room temperature but induced well in the second induction when cultured at 37ºC (Figure S1). The transient expression profile can be attributed to the DH5-α background used and the presence of a LVA protease tag which is not present in any of the other strains tested.

Using the same approach we tested nine engineered arsenic responsive bacterial strains^20,21^. We chose arsenic responsive strains for use in a biodisplay for low-cost, continuous environmental arsenic monitoring. After overnight growth, we flowed LB medium supplemented with 0 μg/l, 10 μg/l, 100 μg/l and 500 μg/l sodium-arsenite for 24 hours and quantitated GFP expression over time. The experiment was performed at room temperature. Strain EDI_as_4 in particular showed high expression of GFP and detected sodium-arsenite down to 10 μg/l (Figure S2). We thus chose this strain for all further arsenic sensing applications.

This series of experiments demonstrated that our biodisplay is a viable high-throughput platform for longterm culturing and characterization of bacterial strains. Theoretically, up to 768 strains can be characterized in parallel on this platform under dynamically changing media conditions^15^, which is otherwise difficult to achieve using existing standard microplate batch cultures or low-throughput microfluidic devices.

### Biodisplay

High levels of arsenic in groundwater are common in a number of countries including China, India, and the USA. Not only is arsenic a concern in water supplies, high arsenic levels have also been found in a number of food sources^22^, particularly rice^23,24^. In Bangladesh, an estimated 20 – 45 million people are affected by arsenic levels that exceed the national standard of 50 µg/l and the WHO / EPA standard of 10 µg/l^25^. Worldwide it is estimated that over 200 million people are exposed to unsafe arsenic levels of 10 µg/l or above^26^. Arsenic also tops the ATSDR substance priority list^27^, which lists those substances “determined to pose the most significant and potential threat to human health”. Our biodisplay could serve as a continuous environmental water monitor that is low-cost, deployable in resource limited settings, and easy to interpret.

We programmed our biodisplay with an arsenic sensing *E. coli* strain (EDI_as_4) spelling “As”, the elemental symbol for arsenic, and a “skull and crossbones” symbol commonly used to warn of toxic substances (Figure 3a). All pixels not programmed with the arsenic sensing strain were programmed with a non-fluorescent strain to prevent invasion and cross-contamination by the arsenic sensing strain. After overnight growth, sodium-arsenite in tap water was sampled every 75 minutes by the device using the sampling routine described above. The biodisplay was kept at room temperature for the entire duration of the sampling routine. Using the research grade fluorescent microscope for readout, the biodisplay detected as little as 10 µg/l sodium-arsenite, and 20 µg/l sodium-arsenite was detected with the handheld low-cost fluorescent USB microscope (Figure S3). Fluorescent signal and the appearance of the “As” and “skull and crossbones” symbols was detected after 10 hours by the research grade microscope (Figure S4). After 24 hours a strong fluorescent signal had developed that could be easily imaged with a low-cost handheld USB microscope. A negative control biodisplay, which was run under identical conditions but exposed to tap water without added arsenic showed no detectable fluorescent signal. We also performed an experiment demonstrating that the biodisplay is able to sense arsenic containing water after 1 day of continuous operation. After overnight growth, arsenic-free tap water was sensed by the biodisplay for 24 hours. After the 24 hour period, the arsenic-free water source was replaced with water containing 50 µg/l sodium-arsenite and an increase in fluorescent signal was observed over time (Figure S5).

**Figure 3.**
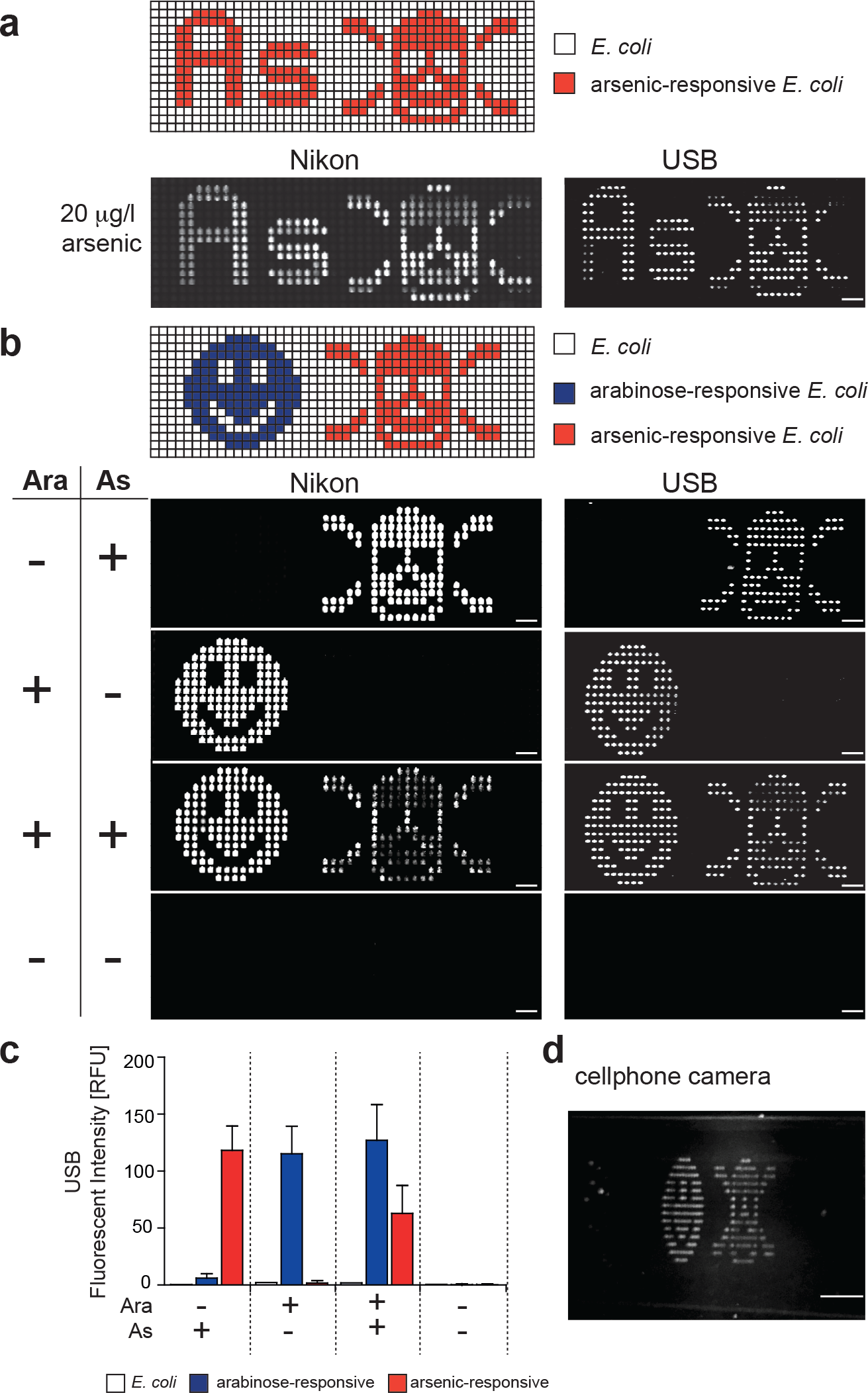
(**a**) Arsenic-responsive *E. coli* was spotted according to the indicated pattern, spelling “As” and forming a “skull and crossbones” symbol. 20 µg/l of sodium-arsenite in tap water was sampled and images were acquired using an epifluorescent and USB microscope after 24 hours. (**b**) Multiplexed detection of arabinose (blue) and arsenic (red) by two *E. coli* strains that were spotted to display a “smiley” and a “skull and crossbones” symbol, respectively. Epifluorescent and USB microscope images of the chip were acquired after flowing all combinations of 6.66 mM arabinose and 100 µg/l sodium-arsenite in tap water in four independent experiments. Scale bars: 750 µm (**c**) Quantitation of GFP intensity after 24 hours of induction, measured using the USB microscope. (**d**) The device exposed to both arsenic and arabinose, was imaged using a mobile phone. The image was obtained using the illumination with blue LEDs and a band-pass filter, placed in front of the cellphone camera. Scale bar: 3.5 mm.

Given the large number of programmable pixels and high pixel density on our biodisplay we explored the possibility of generating a biodisplay that could sense and report the presence or absence of multiple substances in a sample. We selected the same arsenic sensing strain as above and the iGEM GFP_3 strain for arabinose sensing. The arsenic sensing strain was spotted in the form of a “skull and crossbones” pattern whereas the arabinose sensing strain was spotted in a “smiley” pattern. We performed four separate experiments exposing a new biodisplay to one of the four possible combinations of arsenic and arabinose. After sampling for 24 hours, the biodisplay accurately reported the presence / absence of these two small molecules (Figure 3b). We note that the arsenic-sensing strain used (EDI_as_4) was engineered to be tunable by arabinose, explaining the difference in intensity of the “skull and crossbones” symbol between experiments sampling arsenic and arabinose versus arsenic alone^20^. The patterns could be readily imaged using a research grade and USB fluorescent microscope. Because of the ease with which the symbols could be visualized we also tested whether the device could be imaged directly with a cellphone using a blue LED flashlight for illumination and a fluorescence emission filter held in front of the cellphone camera. Both symbols could readily be imaged with this simple setup (Figure 3d). The symbols could also be seen by eye using the LED flashlight and emission filter. Imaging was performed in an illuminated laboratory during daylight hours, with a small curtain shielding the device from direct sunlight (Figure 3d, Figure S6).

### Spores enable long-term storage

In order to develop a biodisplay that can withstand long-term storage and shipping without requiring a cold chain or special conditions, we explored the use of bacterial spores. Bacterial spores are well known for their incredible robustness to adverse environments^28,29^ and thus would be ideal for programming a long-term storage biodisplay.

We generated spores from two *B. subtilis* strains, each containing a genomically integrated P*hy-spank* promoter driving the expression of either mCherry or GFP. The spores were spotted onto glass slides together with *B. subtilis* and *E. coli* cells. The slides were aligned and bonded to a microfluidic device. These biodisplays were then stored for one day or one month at 40ºC and 80ºC. After the storage period we introduced LB medium (Figure 4a,b). Neither *E. coli* nor *B. subtilis* cells could be grown after any of the storage condition tested. Only when spotted in the presence of glycerol and without an extended storage period could *E. coli* and *B. subtilis* cells be grown on-chip. *B. subtilis* spores, on the other hand, germinated after all storage conditions, notably even after incubation at 80ºC for one month. Germination was observed as early as 1.5 hours after the start of the culturing routine. To test whether a spore-based biodisplay retains sensing capability after storage at 80ºC for one month, we induced reporter expression with 1 µM IPTG and observed a rapid increase of fluorescence within 90 minutes. *B. subtilis* biopixels not exposed to IPTG remained non-fluorescent (Figure 4c). To decrease the risk of environmental biological and chemical release, the entire spore biodisplay with attached tygon tubing was embedded in PDMS (Figure S7a). No adverse effects on spore germination and induction were observed (Figure S7b,c). Increasing the flow pressure from 20 kPa to 140 kPa as well as the actuation pressure of the control valves from ~70 kPa to 172 kPa did not lead to device failure, demonstrating its improved tolerance to higher pressures.

**Figure 4.**
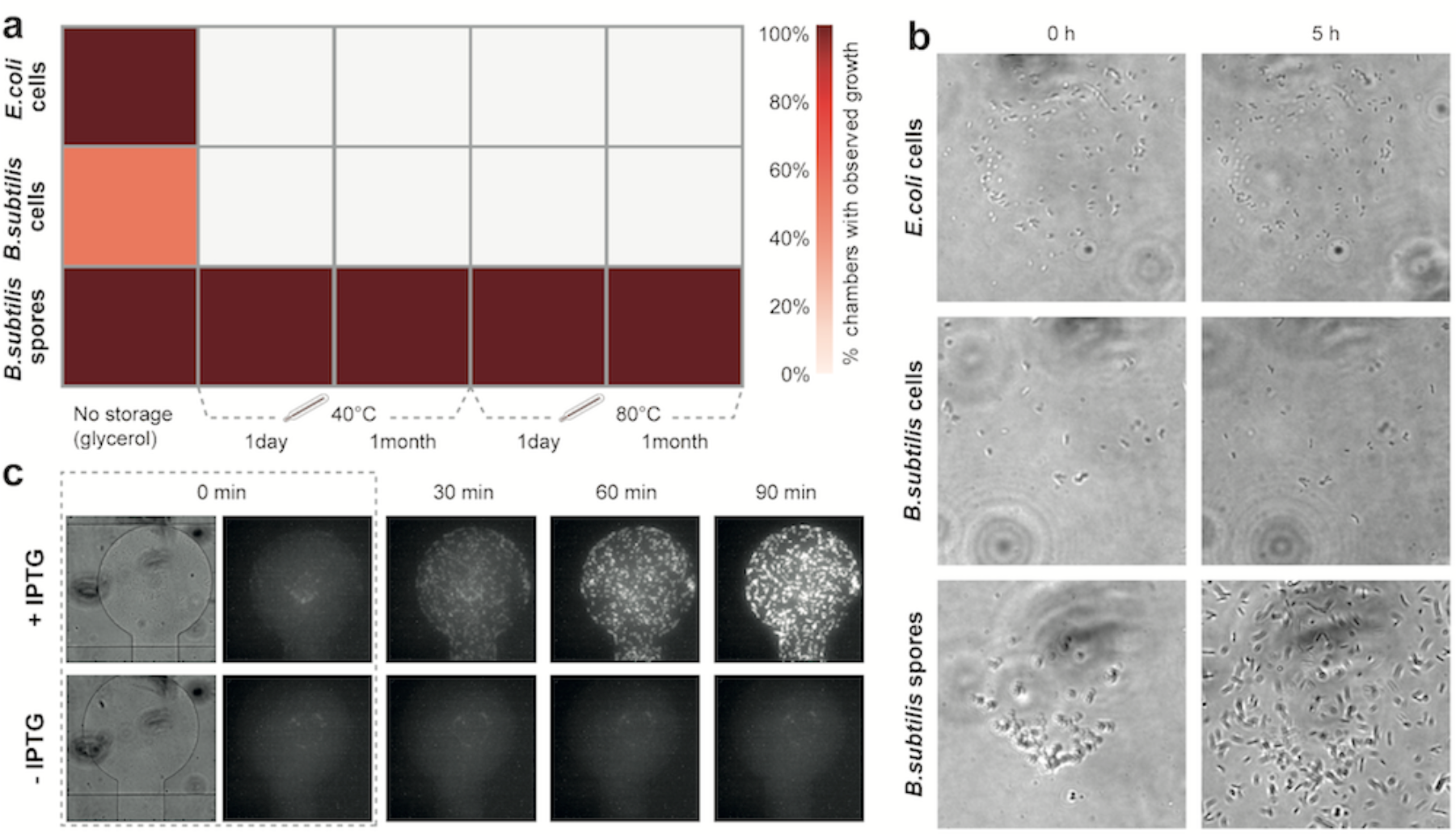
**(a)** Comparison of *E.coli* cell, *B.subtilis* cell and *B.subtilis* spore recovery under different storage conditions. Each table entry represents the number of chambers with observed growth divided by the total number of spotted chambers for a specific temperature and time condition. **(b)** Brightfield images of a magnified view of the central part of representative pixels after storage at 80°C for 1 month. Images were acquired immediately after spots were resuspended in medium and after a 5 hour incubation at 37°C. **(c)** Time-lapse images of germinated P*hy-spank*-mCherry *B. subtilis* spores after storage at 80ºC for one month. Cells were induced with 1 µM IPTG.

### Summary

We developed a bio-hardware device by combining engineered bacterial strains with a microfluidic chip operated under the control of an automated system. The resulting biodisplay consists of 768 independently programmable biopixels and we showed that it could culture bacterial strains for extended periods of time. Continuous culturing with the capability of dynamically changing media makes the biodisplay a useful platform for high-throughput bacteria cell analysis and may find application in high-throughput screening and characterization of synthetically engineered bacterial strains.

Biological systems are powerful sensors, but challenging to deploy for several reasons. First, engineered biological systems generally need to be cultivated under fairly well controlled conditions, and often require the presence of antibiotics in order to maintain the synthetic network. Second, a major concern limiting the applicability of synthetic systems is the possible escape of genetically modified cells into the environment. Not only is the escape of GMOs a concern, but also the release of genetic material, which could be taken up by environmental bacteria through horizontal gene transfer. Our biodisplay solves both issues by providing a platform that enables long-term culturing, while decreasing the risk of release of genetic and chemical material into the environment. The small scale of the device requires low quantities of medium and antibiotics: ~2 ml for a 6 day experiment. The device outflow is sterilized and collected to prevent environmental contamination with biological and chemical components.

The ability to culture bacterial sensors on the biodisplay for extended periods of time with frequent readout makes the biodisplay a potentially useful environmental monitoring tool. We showed that our biodisplay can detect arabinose and sodium-arsenite in tap water using bacterial strains previously engineered by other research groups^20,30^ and student teams^31^. Several heavy metal sensors have already been engineered, including arsenic^32,33^, mercury^33^, and lead sensors^34^. Information processing systems also exist, including genetic toggle switches^35^, logic functions^36^, band-pass filters^37^, and event counters^38^. By combining event counters or toggle switches with sensors it is possible to implement memory, which could be useful in instances when the biodisplay can be monitored only intermittently, but a transient contamination should be logged and reported. The use of band-pass filters would enable the development of a biodisplay that reports the concentration of an analyte without requiring quantitative analysis of the reported signal. The biodisplay can also reduce the complexity of engineered strains, since functions can potentially be distributed amongst several strains^39^. For example, to generate a sensor that senses and reports the presence/absence of three different substances wouldn’t require incorporation of all three functions into a single strain but could instead be implemented in three individual strains. It may also be possible to develop chemiluminescent biodisplays or color biodisplays using chromogenic proteins. Communication between pixels as previously demonstrated using hydrogen peroxide^17^ could be used to couple sensing biopixels, logic biopixels, and reporter biopixels.

Unlike previous biopixel arrays^17^ where each pixel contained the same bacterial strain, each pixel on our display can be specifically programmed with a different strain, drastically increasing the multiplexing capacity and information content of our biodisplay with a maximum of 768 unique strains, each of which could contain multiple, genetically encoded functions. A previous demonstration of integrating an *E. coli* arsenic biosensor in a microfluidic device was limited to a single culture, employed a complex scheme that decoupled culturing from measurement, and used a research-grade fluorescent microscope for readout ^40^. A scheme called InfoBiology was used for the transmission of information using arrays of bacterial cells spotted on microtiter sized agar plates^41^. 144 colonies of a handful of strains were spotted in this scheme, but lacked integration with a microfluidic device for cell culturing and environmental sampling.

We previously developed a low-cost, portable hardware system that contains all necessary components for microfluidic device operation, making the biodisplay readily deployable in resource limited settings^42^. By using bacterial spores the biodisplay can be stored and shipped under ambient conditions and is viable for extended periods of time. The material cost of the biodisplay itself is negligible at less than 1 USD per device. In this particular example a bio-hardware platform applied to continuous environmental monitoring of arsenic levels in water could replace a USD 26’000 instrument with an annual running cost of over USD 3’000 (Ova 7000, Modern Water Monitoring Limited). The relatively slow response time of our biodisplay is acceptable in this and other applications, where continued chronic exposure of a substance is to be avoided. For time critical applications the response time of the biosensors would need to be improved, for example by developing phosphorylation based sensors and reporters rather than transcriptional based genetic networks^43^.

Programmable biopixel displays drastically simplify device readout and interpretation. Instead of reporting the presence of arsenic using a display-wide pattern of signal oscillations^17^, our biodisplay generates an easy to interpret “skull and crossbones” symbol. We showed that this concept can be easily multiplexed using different symbols to report the presence of multiple small molecules. The number of biopixels is scalable with previous work on two layer polydimethylsiloxane (PDMS) chips having demonstrated up to 4160 unit cells per device^44,45^. Immediate optical readout without specialized equipment combined with the fact that computing and decision-making steps are conducted by the biodisplay itself make it low-cost, easy to deploy, and easy to interpret. Bio-hardware hybrid devices therefore combine the advantages of biological systems with the advantages of mechanical, electrical and optical systems. The ability to engineer biological systems on the molecular level and combine them with novel hardware opens new opportunities for how biological, mechanical, electrical and optical systems are integrated and the types of functions they can perform.

## METHODS

### Materials

All bacterial strains and plasmids used in the study are listed in Table S1. Cells for arsenic detection were a gift from Baojun Wang (University of Edinburgh): pBW103ParsR-Amp30C (Addgene plasmid # 78638), pBW300ParsR-Amp32T (Addgene plasmid # 78652), pBW102ParsR-Amp32C (Addgene plasmid # 78637) and pBW101ParsR-gfp (Addgene plasmid # 78636). LB medium was purchased from Applichem Panreac (A0954). L-(+)-Arabinose (A3256-100G), 0.05M (35000-1L-R) and chloramphenicol (23275) were purchase from Sigma. Kanamycin sulfate (T832.1) was bought from Carl Roth. Isopropyl-*β*-D-thiogalactoside for spore induction was purchased from Roche. Schaeffer and Fulton Spore Stain Kit was bought from Sigma Aldrich.

### Device fabrication

Device molds were made using standard photolithography techniques. The mold for the control layer was made using GM 1070 SU-8 photoresist (Gersteltec Sarl, Switzerland) with a height of 30 µm. The flow layer was made with AZ9260 photoresist (Gersteltec Sarl, Switzerland) with a thickness of 14 µm. A Suss MJB4 single side mask aligner was used to expose the wafers. The flow mold was baked for 2 hours at 135°C to anneal the flow channels. The molds were then treated with chlorodimethylsilane (DMCS) prior to being exposed to PDMS (Sylgard 184). The double layer microfluidic device were made by using multilayer soft lithography^46^. For the flow layer, PDMS was prepared with a ratio of 20:1 (part A:B) and spin coated with a speed of 3000 rpm for 1 minute, in order to get a thickness in the range of 20 – 40 μm. The control mold was placed in a petri dish and coated with a 5:1 ratio of PDMS, in order to get a ~ 0.5 cm thick device. Both layers were baked for 30 minutes at 80°C. Then the control layer was cut and peeled from the control layer mold and placed on top of the flow layer. Alignment of the two layers was performed by eye using a stereomicroscope; markers, placed around the main structure of the device aided the alignment process. After alignment the two PDMS layers were baked for an additional 1.5 hours at 80°C. The chip was then aligned to a spotted glass slide.

### Cell culture, transformation and spore formation

*E. coli* and *B. subtilis* cells were cultured in LB medium at 37°C and 200 rpm with appropriate antibiotics, when required. All plasmid transformations and *E.coli* studies were performed in DH5-α cells. *B. subtilis* sporulation was performed as described previously^47^. For spore enrichment, a three day old *B. subtilis* cell suspension was centrifuge for 1 min at 15000 rpm, the pellet was washed three times with water. Schaeffer and Fulton spore staining was performed to verify the presence of spores.

### Cell arraying

Overnight cultures of *E. coli* and *B. subtilis* were centrifuged for 5 min at 3000 rpm. For the *E. coli* display the cell pellet was resuspended in 100 µl LB with 10% glycerol and appropriate antibiotics. For the spore biodisplay spores were resuspended in water. Cell suspensions and *B. subtilis* spore solutions were plated in conical, polypropylene 96-well plates. The samples were spotted with a 0.7 nl delivery volume pin (946MP2B Arrayit) on a glass slide by using a microarray robot (QArray2, Genetix). Glass slides were coated with epoxysilane (3-Glycidoxypropyl-dimethoxymethylsilane 97% AC216545000 Acros organic). The spotting parameters were as follows: 100 ms inking time, 10 ms stamping time, max number of stamps per ink 30. A wash procedure was included that consisted of washing the spotting pin with 70% ethanol for 2 s, washing with water for 2 s followed by pin drying. Depending on the spotting pattern and number of samples used the spotting time varied between 10 to 20 minutes.

### Cell-display culturing

The *E. coli* array was aligned to a PDMS chip and incubated for 1 hour at 37°C. The spore-array was aligned to a MITOMI^48^ PDMS chip and incubated depending on the experimental procedure as follows: 40°C for 1 day, 40°C for 1 month, 80°C for 1 day, or 80°C for 1 month. After the incubation step the devices were placed on a temperature-controlled glass plate (H401-NIKON-TI_SR_GLASS/H401_T_CONTROLLER; Okolab) at 37°C. Tygon tubes were filled with deionized water and connected to the inlets of the control lines, pressure was applied in order to actuate the valves. For the *E. coli* display 68.9 kPa and 13.8 kPa were used, for the control and flow lines respectively. For the spore display 103.4 kPa and 24.1-27.5 kPa were used instead. For the spore display we used a 768 unit cell MITOMI chip^48^, but performed the same culturing routine as on the biodisplay device. For the cell culturing and sampling routines a custom-written LabVIEW program was used. During the culturing, LB medium with appropriate antibiotics and lysis buffer containing 30 mM of NaOH (06203-1KG Sigma Aldrich) and 12% SDS (L3771-100G Sigma) was used. Depending on the experimental procedure LB medium or tap water was supplemented with arabinose, sodium-arsenite, or IPTG. The level of arsenic in the tap water was measured by a colorimetric arsenic test kit (MQuant™ Arsenic Test, Sigma Aldrich) with a LOD of 5 µg/l. Arsenic levels in our tap water were below the detection limit, and based on publically available data are likely below 2 µg/l^49^.

### Imaging

Image acquisition was performed on a Nikon ECLIPSE *Ti* automated microscope equipped with a LED Fluorescence Excitation System and a Hamamatsu ORCA-Flash 4.0 camera controlled by NIS Elements. Images were taken at 40x magnification (SPlan Fluor, ELWD 40x/0.60, ∞/0.2, WD 3.6-2.8, Nikon) in fluorescent and bright field mode. Images of all biopixel units were stitched together using either the Grid/Collection Stitching plugin in Fiji or a custom-written Python script. Fluorescent measurements were performed using Genepix software. A FITC USB fluorescent microscope (AM4113T-GFBW, Dino-Lite) was used to acquire fluorescent images. Images of 9 sub-sections of the device were taken, using lower magnification (10x), and stitched. Pictures were also taken using a cellphone camera. A band-pass filter, centered at 530 nm with a 40 nm bandwidth, was placed in front of the camera of the mobile phone, and the LEDs of the FITC USB microscope were used for illumination of the biodisplay.

## SUPPORTING INFORMATION

**The Supporting Information document includes:**

**Figure S1.** Biodisplay for strain characterization at 37°C.

**Figure S2.** Arsenic-responsive *E. coli.*

**Figure S3.** Arsenic biodisplay.

**Figure S4.** Arsenic biodisplay over time.

**Figure S5.** Delayed arsenite sensing.

**Figure S6.** Cellphone image acquisition.

**Figure S7.** Embedded spore biodisplay.

**Table S1.** List of plasmids and strains used in this work.

## ACKNOWLEDGEMENTS

The work was supported by the Ecole Polytechnique Federale de Lausanne and a Swiss National Science Foundation grant (CR23I2 140697). W
e thank Pernille Rainer and Kilian Cochet for the initial characterization of cell spotting, Philippe Lenzen for the preliminary characterization of the spore-display and Nadanai Laohakunakorn and Mathieu Quinodoz for help with LabVIEW programming. We also thank Professor Baojun Wang (University of Edinburg) and Professor Jan Roelof van der Meer (University of Lausanne) for the kind gift of arsenic-responsive *E. coli* strains.

## ATUHOR CONTRIBUTIONS

F.V., E.P. and S.J.M. designed research, analyzed the data and wrote the manuscript. F.V. performed the microfabrication and F.V. and E.P. performed the experiments.

## REFERENCES

(1) van der Meer, J. R., and Belkin, S. (2010) Where microbiology meets microengineering: design and applications of reporter bacteria. Nat. Rev. Microbiol. 8, 511–522.

(2) Bereza-Malcolm, L. T., Mann, G., and Franks, A. E. (2015) Environmental Sensing of Heavy Metals Through Whole Cell Microbial Biosensors: A Synthetic Biology Approach. ACS Synth. Biol. 4, 535–546.

(3) Szita, N., Polizzi, K., Jaccard, N., and Baganz, F. (2010) Microfluidic approaches for systems and synthetic biology. Curr. Opin. Biotechnol. 21, 517–523.

(4) Gulati, S., Rouilly, V., Niu, X., Chappell, J., Kitney, R. I., Edel, J. B., Freemont, P. S., and deMello, A. J. (2009) Opportunities for microfluidic technologies in synthetic biology. J. R. Soc. Interface 6, S493–S506.

(5) Wright, O., Stan, G.-B., and Ellis, T. (2013) Building-in biosafety for synthetic biology. Microbiol. Read. Engl. 159, 1221–1235.

(6) Whitesides, G. M. (2006) The origins and the future of microfluidics. Nature 442, 368–373.

(7) Maerkl, S. J. (2009) Integration column: Microfluidic high-throughput screening. Integr. Biol. Quant. Biosci. Nano Macro 1, 19–29.

(8) Nobs, J.-B., and Maerkl, S. J. (2014) Long-term single cell analysis of S. pombe on a microfluidic microchemostat array. PloS One 9, e93466.

(9) Bennett, M. R., and Hasty, J. (2009) Microfluidic devices for measuring gene network dynamics in single cells. Nat. Rev. Genet. 10, 628–638.

(10) Garcia-Cordero, J. L., and Maerkl, S. J. (2013) A 1024-sample serum analyzer chip for cancer diagnostics. Lab. Chip 14, 2642–2650.

(11) Garcia-Cordero, J. L., and Maerkl, S. J. (2016) Mechanically Induced Trapping of Molecular Interactions and Its Applications. J. Lab. Autom. 21, 356–367.

(12) Morin, S. A., Shepherd, R. F., Kwok, S. W., Stokes, A. A., Nemiroski, A., and Whitesides, G. M. (2012) Camouflage and display for soft machines. Science 337, 828–832.

(13) Riedel-Kruse, I. H., Chung, A. M., Dura, B., Hamilton, A. L., Lee, B. C. (2011) Design, engineering and utility of biotic games. Lab Chip 11, 14–22.

(14) Liu, X., Tang, T.-C., Tham, E., Yuk, H., and Lin, S. (2017) Stretchable Living Materials and Devices with Hydrogel-Elastomer Hybrids Hosting Programmed Cells. Proc. Natl. Acad. Sci. 114, 2200–2205.

(15) Dénervaud, N., Becker, J., Delgado-Gonzalo, R., Damay, P., Rajkumar, A. S., Unser, M., Shore, D., Naef, F., and Maerkl, S. J. (2013) A chemostat array enables the spatio-temporal analysis of the yeast proteome. Proc. Natl. Acad. Sci. U. S. A. 110, 15842–15847.

(16) Rajkumar, A. S., Dénervaud, N., and Maerkl, S. J. (2013) Mapping the fine structure of a eukaryotic promoter input-output function. Nat. Genet. 45, 1207–1215.

(17) Prindle, A., Samayoa, P., Razinkov, I., Danino, T., Tsimring, L. S., and Hasty, J. (2011) A sensing array of radically coupled genetic “biopixels.” Nature 481, 39–44.

(18) Hansen, C. L., Skordalakes, E., Berger, J. M., and Quake, S. R. (2002) A robust and scalable microfluidic metering method that allows protein crystal growth by free interface diffusion. Proc. Natl. Acad. Sci. U. S. A. 99, 16531–16536.

(19) Balagaddé, F. K., You, L., Hansen, C. L., Arnold, F. H., and Quake, S. R. (2005) Long-term monitoring of bacteria undergoing programmed population control in a microchemostat. Science 309, 137–140.

(20) Wang, B., Barahona, M., and Buck, M. (2014) Engineering modular and tunable genetic amplifiers for scaling transcriptional signals in cascaded gene networks. Nucleic Acids Res. 42, 9484–9492.

(21) Merulla, D., Hatzimanikatis, V., and van der Meer, J. R. (2013) Tunable reporter signal production in feedback-uncoupled arsenic bioreporters. Microb. Biotechnol. 6, 503–514.

(22) Lynch, H. N., Greenberg, G. I., Pollock, M. C., and Lewis, A. S. (2014) A comprehensive evaluation of inorganic arsenic in food and considerations for dietary intake analyses. Sci. Total Environ. 496, 299–313.

(23) Hojsak, I., Braegger, C., Bronsky, J., Campoy, C., Colomb, V., Decsi, T., Domellöf, M., Fewtrell, M., Mis, N. F., Mihatsch, W., Molgaard, C., and van Goudoever, J. (2015) Arsenic in Rice. J. Pediatr. Gastroenterol. Nutr. 60, 142–145.

(24) Meharg, A. A., Deacon, C., Campbell, R. C. J., Carey, A.-M., Williams, P. N., Feldmann, J., and Raab, A. (2008) Inorganic arsenic levels in rice milk exceed EU and US drinking water standards. J. Environ. Monit. 10, 428–431.

(25) (2016) WHO | Arsenic. WHO.

(26) Naujokas, M. F., Anderson, B., Ahsan, H., Aposhian, H. V., Graziano, J. H., Thompson, C., and Suk, W. A. (2013) The broad scope of health effects from chronic arsenic exposure: update on a worldwide public health problem. Environ. Health Perspect. 121, 295–302.

(27) Priority List of Hazardous Substances | ATSDR.

(28) Setlow, P. (2006) Spores of Bacillus subtilis: their resistance to and killing by radiation, heat and chemicals. J. Appl. Microbiol. 101, 514–525.

(29) Atrih, A., and Foster, S. J. (1999) The role of peptidoglycan structure and structural dynamics during endospore dormancy and germination. Antonie Van Leeuwenhoek 75, 299–307.

(30) Merulla, D., Hatzimanikatis, V., and van der Meer, J. R. (2013) Tunable reporter signal production in feedback-uncoupled arsenic bioreporters. Microb. Biotechnol. 6, 503–514.

(31) parts.igem.org.

(32) Kaur, H., Kumar, R., Babu, J. N., and Mittal, S. (2015) Advances in arsenic biosensor development – A comprehensive review. Biosens. Bioelectron. 63, 533–545.

(33) Wright, O., Delmans, M., Stan, G.-B., and Ellis, T. (2015) GeneGuard: A Modular Plasmid System Designed for Biosafety. ACS Synth. Biol. 4, 307–316.

(34) Bereza-Malcolm, L., Aracic, S., and Franks, A. E. (2016) Development and Application of a Synthetically-Derived Lead Biosensor Construct for Use in Gram-Negative Bacteria. Sensors 16, E2174.

(35) Gardner, T. S., Cantor, C. R., and Collins, J. J. (2000) Construction of a genetic toggle switch in Escherichia coli. Nature 403, 339–42.

(36) Nielsen, A. A. K., Der, B. S., Shin, J., Vaidyanathan, P., Paralanov, V., Strychalski, E. A., Ross, D., Densmore, D., and Voigt, C. A. (2016) Genetic circuit design automation. Science 352, aac7341–aac7341.

(37) Basu, S., Gerchman, Y., Collins, C. H., Arnold, F. H., and Weiss, R. (2005) A synthetic multicellular system for programmed pattern formation. Nature 434, 1130–1134.

(38) Friedland, A. E., Lu, T. K., Wang, X., Shi, D., Church, G., and Collins, J. J. (2009) Synthetic Gene Networks That Count. Science 324, 1199–1202.

(39) Shong, J., Rafael, M., Diaz, J., and Collins, C. H. (2012) Towards synthetic microbial consortia for bioprocessing. Curr. Opin. Biotechnol. 23, 798–802.

(40) Buffi, N., Beggah, S., Truffer, F., Geiser, M., van Lintel, H., Renaud, P., van der Meer, J. R., Geiser, M., Buttgenbach, S., Franco-Lara, E., and Krull, R. (2016) An automated microreactor for semi-continuous biosensor measurements. Lab Chip 16, 1383–1392.

(41) Palacios, M. A., Benito-Peña, E., Manesse, M., Mazzeo, A. D., Lafratta, C. N., Whitesides, G. M., and Walt, D. R. (2011) InfoBiology by printed arrays of microorganism colonies for timed and on-demand release of messages. Proc. Natl. Acad. Sci. U. S. A. 108, 16510–16514.

(42) Piraino, F., Volpetti, F., Watson, C., and Maerkl, S. J. (2016) A Digital–Analog Microfluidic Platform for Patient-Centric Multiplexed Biomarker Diagnostics of Ultralow Volume Samples. ACS Nano 10, 1699–1710.

(43) Purnick, P. E. M., and Weiss, R. (2009) The second wave of synthetic biology: from modules to systems. Nat. Rev. Mol. Cell Biol. 10, 410–422.

(44) Fordyce, P. M., Gerber, D., Tran, D., Zheng, J., Li, H., DeRisi, J. L., and Quake, S. R. (2010) De novo identification and biophysical characterization of transcription-factor binding sites with microfluidic affinity analysis. Nat. Biotechnol. 28, 970–5.

(45) Garcia-Cordero, J. L., and Maerkl, S. J. (2016) Mechanically Induced Trapping of Molecular Interactions and Its Applications. J. Lab. Autom. 21, 356–367.

(46) Unger, M. A., Chou, H.-P., Thorsen, T., Scherer, A., and Quake, S. R. (2000) Monolithic Microfabricated Valves and Pumps by Multilayer Soft Lithography. Science 288, 113–116.

(47) Harwood, C. R., and Cutting, S. M. (1990) Molecular biological methods for Bacillus. Wiley, Chichester; New York.

(48) Maerkl, S. J., and Quake, S. R. (2007) Systems approach to measuring the binding energy landscapes of transcription factors. Science 315, 233–237.

(49) Pfeifer, Hassouna M., Plata, N., H. R. (2012) Arsenic in the different environmental compartments of Switzerland: an updated inventory., in 4th International Conference on Metals and Related Substances in Drinking Water (METEAU).

